# Partial Activation of PPAR-γ by Synthesized Quercetin Derivatives Modulates TGF-β1-Induced EMT in Lung Cancer Cells

**DOI:** 10.1101/2022.11.23.517612

**Authors:** Sangeeta Ballav, Amit Ranjan, Soumya Basu

**Affiliations:** Cancer and Translational Research Laboratory, Dr. D.Y. Patil Biotechnology and Bioinformatics Institute, Dr. D.Y. Patil Vidyapeeth, Tathawade, Pune, Maharashtra – 411 033, India

**Keywords:** Epithelial–mesenchymal transition, Quercetin, Partial agonist, PPAR-γ, TGF-β1, Lung cancer

## Abstract

Non-small cell lung cancer (NSCLC) possess very low survival rate due to poor response to chemotherapy and late detection. Epithelial to mesenchymal transition (EMT) is regarded as a major contributor to drive metastasis during NSCLC progression. Towards this, transforming growth factor-beta 1 (TGF-β1) is the key driver that endows cancer cells with increased aggressiveness. Recently, our group synthesized a series of Schiff base quercetin derivatives (QDs) and ascertained their effectiveness on EMT markers of A549 cell line. Our study evidenced that EMT process was counteracted via the partial activation of a nuclear hormone receptor, Peroxisome proliferator-activated receptor (PPAR)-γ through QDs. Hence, here we extended our work to investigate the interplay between PPAR-γ partial activation by synthesized QDs, TGF-β1-induced EMT and migration in human lung cancer A549 cells. The results revealed that TGF-β1 played a critical role in suppressing PPAR-γ, which was markedly reversed and increased by partial agonists; QUE2FH and QUESH at both protein and transcriptional level. Compared to full agonists, rosiglitazone could not elevate PPAR-γ expression in the presence of TGF-β1 and had negligible effect on translocation of PPAR-γ to nucleus. The partial agonists not only stimulated PPAR-γ in balanced manner but also prevented the loss of E-cadherin and acquisition of TGF-β1-induced mesenchymal markers (Snail, Slug, Vimentin and Zeb-1). Subsequently, the effects were accompanied by attenuation of TGF-β1-induced migratory ability of A549 cells. Together, with the balanced activation profile of PPAR-γ ligands, our findings suggest that these novel partial agonists may serve as potential anti-cancer agents to impede metastasis.

## Introduction

Non-small cell lung carcinoma (NSCLC) being a debilitating disease, addresses the changes occurring during the course of tumour progression and one of the major hallmark of such changes is known to be epithelial–mesenchymal transition (EMT). EMT is characterized by transition of polarized epithelial cells to motile mesenchymal cells, leading to manifested biological morphogenetic process [1]. Transforming growth factor beta 1 (TGF-β1), a multifunctional cytokine, is a key player to have greater impact on EMT that deliberately contributes to a complex milieu, eventually causes the tumour microenvironment to develop metastasis and therapy resistance of tumours [2]. PPAR-γ has become the lime light in the field of drug discovery and development since they have tuned for its unique and attractive therapeutic intervention [3-5]. Rekha and colleagues has illustrated in their study about the activation of PPAR-γ independent of Smad phosphorylation inhibition by two thiazolidinediones (TZDs); rosiglitazone and ciglitazone in lung cancer cells. They also found the inhibition of mesenchymal markers and matrix metalloproteases by siRNA-mediated knockdown of PPAR-γ which authenticated the action of PPAR-γ to be ligand-dependent [3].

Transforming growth factor-beta 1 (TGF-β1) is well-known for increasing the expression of transcriptional factors of mesenchymal markers and decreasing the E-cadherin transcription [6]. These transcriptional repressors of E-cadherin are needed during Epithelial–mesenchymal transition (EMT) development, suggesting that upregulating E-cadherin expression by PPAR-γ ligands might be a reliable strategy for hindering the gain of mesenchymal markers and subsequent functional phenotype of increased motility during EMT. PPAR-γ is considered as versatile transcription factor that is activated by ligands and exerts anti-proliferative and anti-metastatic effects by modulating TGF-β1-induced EMT pathway [4, 7]. It is believed that ligand serve as potent activator for PPAR-γ, thereby playing a significant role in exerting anti-cancer responses. Although several endogenous ligands have been into the account of triggering PPAR-γ but they seem to exert low affinity [8]. On the other hand, synthetic ligands appear to be promising modality for PPAR-γ, namely, thiazolidinediones (TZDs) and non-steroidal anti-inflammatory drugs (NSAIDs);which act as full agonists leading to an over-activation process [9] that causes deteriorating adverse effects such as higher risk of the cardiovascular system such as heart strokes and congestive heart failure [10], edema, water retention and osteoporosis [11]. Hence, the application has been restricted due to their toxicity profile.

Recently, our group synthesized a series of Schiff base derivatives of quercetin and ascertained their effectiveness on EMT markers ofA549 cell line. Our study evidenced that EMT process was counteracted via the activation of PPAR-γ through the QDs. Hence, in the present study, we have strategized to modulate EMT pathway induced by TGF-β1 by the means of partial activation of PPAR-γ with our novel derivatives of quercetin. As TGF-β1 is a crucial player to have a greater influence on EMT and having verified with PPAR-γ activation by QDs, here we extended our work to investigate the interplay between PPAR-γ, TGF-β1and synthesized QDs. To the best of our knowledge, this is the first time to show inhibition of TGF-β1-induced EMT in lung cancer cells via PPAR-γ partial activation by chemically synthesized small molecules. The compounds that stimulate PPAR-γ in desired manner i.e., lesser effect than synthetic agonists and greater effect than weak agonists are referred to as partial agonist. Various potential endogeneous ligands such as long chain polyunsaturated fatty acids, J-series of prostaglandins and arachidonic acids activate PPAR-γ, but with relatively low affinity, i.e. they act as weak agonists [8]. While the synthetic agonists such as TZD class of drugs serve as full agonists. It is imperative to note that TZDs has reduced the incidence on various cancers [11], however, to date; there is no such evidence for their potential impact for active malignant disease. In addition, owing to their toxicity concerns the administration to cancer patients. Our previous study successfully achieved three partial agonists namely, QUETSC, QUE2FH and QUESH that activated PPAR-γ in balanced manner i.e., lower than RSG (full agonist) and higher than quercetin (weak agonist). Our study focused at ascertaining new insights in the quest of unraveling the high drug-like properties of newly synthesized lead derivatives of quercetin with effective inhibition of EMT induced by TGF-β1on NSCLC cell lines via partial activation of PPAR-γ protein receptor. We believe this strategy can render an astute way to exploit and modulate in the structural design of quercetin.

## Experimental section

### 1. Chemicals and Reagents

Quercetin and Rosiglitazone (RSG) were purchased from Cayman Chemical Company (MI, USA). Roswell Park Memorial Institute Medium (RPMI)-1640 medium, 200 U/mL of penicillin, 0.2 mg/ml of streptomycin and 200mM L-glutamine, fetal bovine serum (FBS), glutamine, 0.25% trypsin/0.038% EDTA, crystal violet stain, 3-(4,5-dimethythiazol2-yl)-2,5-diphenyl tetrazolium bromide (MTT) dye and (4′,6-diamidino-2-phenylindole) DAPI stain were all purchased from HiMedia. TRIzol reagent was purchase from Ambion, life technologies. All the primers used for mRNA expression were obtained from Sigma Aldrich (St. Louis, MO, USA).Enhanced chemiluminescent (ECL) substrate kit was purchased from Bio-rad, USA. Roswell Park Memorial Institute Medium (RPMI)-1640 medium, 200 U/mL of penicillin, 0.2 mg/ml of streptomycin and 200mM L-glutamine, 10% fetal bovine serum (FBS), 1% glutamine (200 mM), 0.25% trypsin/0.038% EDTA, crystal violet stain, 3-(4,5-dimethythiazol2-yl)-2,5-diphenyl tetrazolium bromide (MTT) dye was purchased from HiMedia (Mumbai, India). Enhanced chemiluminescent (ECL) substrate kit was purchased fromBio-rad (CA, USA).Antibodies details are as follows; Monoclonal anti-human antibody: β-actin (mouse, Cloud clone) and polyclonal anti-human antibodies PPAR-γ (rabbit, Cloud clone), EMT markers-anti-Snail (rabbit, Cell Signaling Technology, CA, USA), anti-N-cadherin (rabbit, Cell Signaling Technology, CA, USA), anti-Vimentin (rabbit, Cell Signaling Technology, CA, USA), anti-Slug (rabbit, Cell Signaling Technology, CA, USA), anti-E-cadherin (rabbit, Cell Signaling Technology, CA, USA) and anti-Zeb-1 (rabbit, Cell Signaling Technology, CA, USA). Anti-rabbit and anti-mouse antibodies conjugated to horseradish peroxidase (HRP) were obtained from Cell Signaling Technology, CA, USA.

### 2. Cell culture conditions

The human NSCLC cell lines A549 and NCI-H460 were obtained from the National Centre for Cell Science (NCCS),Pune, Maharashtra, India. Both the cell lines were maintained in RPMI-1640 medium supplemented with 10% FBS, 1% 200 U/mL of penicillin, 1% 0.2 mg/ml of streptomycin, and 1% 200mM L-glutamine in a humidified, 5% CO2, 37°C incubator (Eppendorf, Germany). The media was changed every 2-3 days, and the cells were separated via trypsinization using 0.25% trypsin/0.038% EDTA when they reached 70-80 % confluence.

### 3. Western blot analysis

Total cellular proteins (40µg) from A549 cells were treated with QDs and positive controls,quercetin and RSG for 24hr. Protein concentrations of all the mentioned samples were determined using the Bradford assay.An equal amount ofproteinwasboiled at 95°C for 5 min after the addition of 3X Laemmli buffer, followed by separation on 10% SDS-PAGE and the separated proteins were electrotransferred onto polyvinylidene fluoride (PVDF) membraneusing a Trans-Blot® Turbo™ Transfer System (Bio-Rad Laboratories, Inc., India). The membranes were first blocked with 5% non-fat milk in TBST for 1hr at room temperature, followed by probing with respective primary antibodies following published protocols [12]. The primary antibodies used are as follows; anti-PPAR-γ (1:500), EMT markers-anti-Snail (1:500), anti-N-cadherin (1:500), anti-Vimentin (1:1000), anti-Slug (1:250), anti-E-cadherin (1:500) and anti-Zeb-1 (1:250). Anti-rabbit antibody conjugated to horseradish peroxidase (HRP) (1:1000) was used as the secondary antibody andincubated for 1hr. β-actin was used as a control. The blotswere incubated in ECL Plus reagent, and chemiluminescence was detected on iBright CL1500 Imaging System (Thermo Fischer Scientific) and quantified the density of the bands.

### 4. Preparation of nuclear extracts of treated cells for measurement of PPAR-γ activity

Levels of PPAR-γ and activity of QDs was measured by extracting nuclear fraction of A549 cells using PPAR-γ Transcription Factor Assay Kit (RayBiotech, India). Cells were seeded at a density of 4×10^6^ cells per 90mm tissue culture plate to confluence. Twentyfour-hour post-plating, culture medium was removed and replaced with fresh medium containing the appropriate treatment of QDsand positive controls; quercetin and RSG for 24h. Another set of experiment was carried out by treating the cells with QDs and induced with TGF-β1 (5ng/ml) for 72h. The nuclear extracts were isolated using a NE-PER Nuclear and Cytoplasmic Extraction Reagents (Thermo Scientific), as per the manufacturer’s instructions. The protein concentrations of the resultant supernatants (nuclear extract) were determined using the Bradford method.

### 5. Quantitative measurement of PPAR-γ activity

PPAR-γ activity was assayed in nuclear extracts using the PPAR-γ Transcription Factor Activity Assay Kit (RayBiotech, India), following the manufacturer’s instructions. The bottom of each well is pre-coated with PPAR response element (PPRE) by the manufacturer for capturing free PPAR-γ in samples. The volume of nuclear extracts that was spiked in the assay was adjusted based on protein content of individual samples. Briefly, following blocking of nonspecific binding to the PPRE-coated wells, nuclear extracts or positive control extract (supplied with the kit) were added to the appropriate wells and the plate was incubated for overnight at 4°C with gentle shaking. Primary anti-PPAR-γ was added to each well and the plate was incubated for 1hr at room temperature followed by secondary antibody (anti-rabbit IgG-HRP) addition to each well and incubated for 1hr. The wells were washed with 1X wash buffer provided with the kit for four times after each step. Colorimetric detection of bound antibody was performed by addition of Tetramethylbenzidine (TMB) substrate, incubation at room temperature for 30 min, addition of stop solution and measurement of absorbance at 450 nm using in a microplate spectrophotometer (Epoch Microplate Spectrophotometer, Biotek).

### 6. Transwell migration assay

In our previous study, QUESH was sighted to act hold of wound in A549 cells in scratch assay. Therefore, in the present work the migration of A549 cells were further determined by transwell inserts (Corning Incorporation, New York, NY, USA). Briefly, after relevant treatment with QUESH, quercetin, RSG with or without TGF-β1(5ng/ml), 1 × 10^5^ A549 cells were suspended in 300 μl serum-free RPMI-1640 medium and added into the upper chamber.750 μl complete RPMI-1640 medium with serum was added into the lower chamber. After incubation for 72 h at 37 °C, non-migrating cellsin upper chamber were removed using cotton swab carefully and migratory cells in lower chamber cells were fixed with pre-chilled 4% paraformaldehyde for 5min followed by pre-chilled absolute methanol for 10 min and stained with 0.5% crystal violet at room temperature for 15 min. Representative images were taken under an inverted microscope Magnus Analytical MagVision, Version: x64equipped with a camera (Magnum DC5).

### 7. Statistical analysis

Normally distributed continuous variables are expressed as the means ± standard deviation (S.D.). One-way ANOVA followed by the “Dunnett’s Multiple Comparison Test” method was used to compare the results obtained in different treatment groups. A two-sided P value < 0.05 was considered statistically significant. Statistical analyses were performed using GraphPad Prism version 5.0 software (GraphPad Software, La Jolla, CA, USA).

## Results and Discussion

### 1. TGF-β1 promotes morphological changes in A549 cells at 72h

TGF-β1 is ought to be a potent contributor to EMT by altering the polyglonal appearance of cell adhesion molecules which causes to lose cell-cell contacts, leading to spindle-shaped mesenchymal cells [13]. Studies have well reported that A549 cells undergo changes in their shape from cobble-shaped to elongated fibroblastoid form of mesenchymal characteristics in 72h induction of TGF-β1 [3]. Therefore, we first assessed the morphological difference of A549 cells after inducing them with TGF-β1. As shown in figure 1, we noted a clear difference in the structural appearance of the cells at 72h time incubation. At first, cells were observed to be scattered with cytoskeleton-like extensions and the morphological difference became more evident after 48h. The cells shifted from polygonal (epithelial) to spindle shape (mesenchymal). Hence, this authenticates that TGF-β1 induces EMT in A549 cells within 72 h.

**Figure 1:**
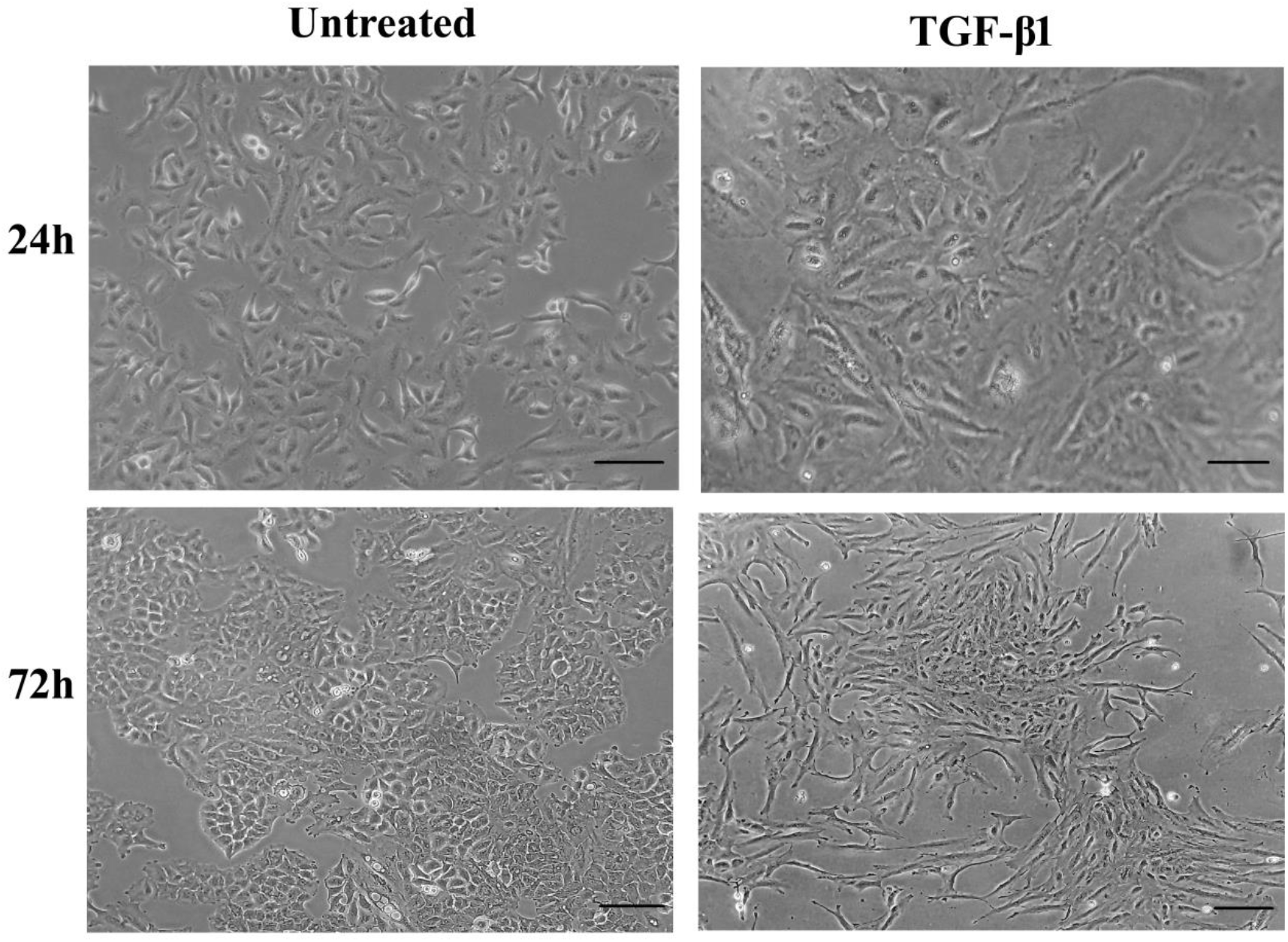
Light microscope analyses: TGF-β1 induces EMT in A549 cells in 72 h. A549 cells were treated with TGF-β1 (5 ng/mL) and the stimulation was monitored for 72h. Representative images are presented at two timelines; 24h and 72h that were captured under Olympus microscope equipped with camera (Magnus Analytical MagVision, Version: x86). Scale bar: 100μm.

### 2. Effect of TGF-β1 on expression of PPAR-γ

As TGF-β1 is a key player to have a greater impact on EMT, we sought to evaluate the probability of TGF-β1 to modulate PPAR-γ expression. In our previous study, we observed the activation of PPAR-γ modulated EMT transition at 24h; therefore we carried out western blot analysis at two intervals; 24h and 72h. According to the quantitative plots (Figure 2), we observed the following pattern: the longer the time exposure given for TGF-β1 incorporation, the greater was the decrease in expression of PPAR-γ. Our analysis reflected that PPAR-γ protein level remarkably decreased after 24 h of TGF-β1stimulation. Hence, statistically significant difference was observed at 72h. This observation is in coherence with other studies where it is speculated that the expression of PPAR-γ is governed by TGF-β1in lung cancer [14]. Hence, the response of TGF-β1for PPAR-γ is time-dependent in nature.

**Figure 2.**
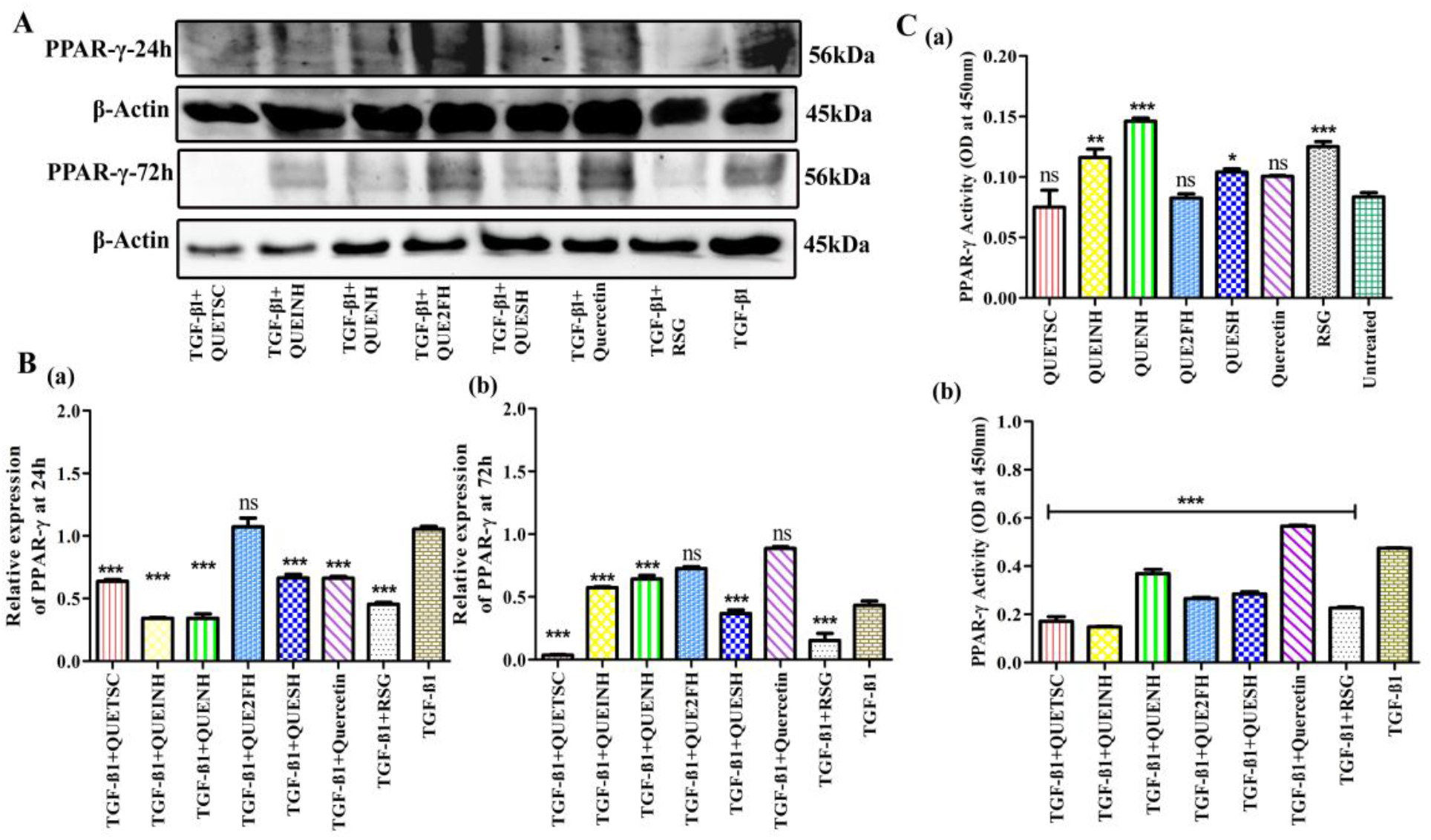
Effect of synthesized QDs on TGF-β1-induced PPAR-γ and its translocation. (A & B) Representative Western blotting images with their relative protein expression level of PPAR-γ at 24h and 72h. (C) PPAR-γ Transcription factor binding activity in nuclear extractions of A549 cells by QDs. (a) Without TGF-β1 (b) with TGF-β1. Each value presented as mean±SD for three replicate determinations for each treatment (p<0.05, one-way ANOVA). The error bar represents SD; *p<0.05, **p<0.01 and ***p < 0.001 compared with control group-TGF-β1.

### 3. Effect of QDs on TGF-β1-induced PPAR-γ

We recently demonstrated thatall the five newly synthesized quercetin derivatives (QDs) functioned as PPAR-γ activatorswhich in turn caused changes in EMT markers in A549 cells. Having verified with PPAR-γ activation by QDs, we next tend to investigate the interplay between PPAR-γ, TGF-β1and synthesized QDs. We evaluated whether the activation of PPAR-γ is affected by QDs on incorporation of TGF-β1 in the cells. As shown in Figure 2 (A), Western blot analysis revealed that the expression of PPAR-γ was diminished after the treatment of TGF-β1. While, the QDs, QUEINH, QUENH, QUE2FH and QUESH increased the expression after 24h treatment. This two-fold elevation in the protein expression was comparable with the parent compound; quercetin which has shown to increase the protein receptor at 72h.In contrast, we noticed a strong suppression in the presence of RSG. This was expected as RSG reports to have full agonist-inducible property and leads to over-activation of PPAR-γ [15]. Thereafter, RSG reversed the action of PPAR-γ in the presence of TGF-β1. These outcomes suggest that QUEINH, QUENH, QUE2FH and QUESH act as PPAR-γ agonist even after incorporation of TGF-β1. However, previously we demonstratedthat QUETSC, QUE2FH, and QUESH led to partial activation of PPAR-γ i.e., lower than RSG (leads to higher expression of PPAR-γ) and higher than quercetin (leads to lower expression of PPAR-γ). Although, after inducing TGF-β1, QUE2FH and QUESH activated the receptor, but QUETSC could not increase PPAR-γ expression. Hence, QUE2FH and QUESH were found to be adhered to its partial activation property. Therefore, these two QDs found to stimulate and reverse PPAR-γ protein expression after inducing TGF-β1 in desired manner i.e., lower than quercetin and higher than RSG.

### 4. QDs promoted nuclear translocation of PPAR-γ

Because PPAR-γ activation solely occurs in nucleus after getting bound with ligand in cytoplasm, we sought to examine the effect of QDs on PPAR-γ transcriptional activity profile with or without TGF-β1. In other words, the assay involves high-throughput mechanism that characterizes protein-ligand interactions by monitoring the functional activity of transcription factors upon binding of agonists to PPAR-γ [16]. At first, we assessed the authenticity of our protocol by determining the expression of PPAR-γ in both cytoplasmic fraction (CF) and nuclear fraction (NF). As shown in Figure S1, PPAR-γ was detected in both cytoplasm and nucleus with substantially more protein content in NF than that of CF. It is obvious that under basal conditions, due to absence of any ligand, expression of PPAR-γ will be seen in cytoplasm too.

Hence, pure translocation exists only in presence of ligands. We have used PCNA as loading control in this experiment (Figure S1) and found three bands of PCNA. Literature review and crystallographic studies suggested that PCNA of human forms a homodimeric or homotrimeric ring that is strongly important for its function and it is found with molecular weights of 30kDa, 60kDa and 90kDa, stating that it forms trimmers [17-18]. It is obvious that under basal conditions, due to absence of any ligand, expression of PPAR-γ will be seen in cytoplasm too. Hence, pure translocation exists only in presence of ligands.

With the treatment of our synthesized PPAR-γ ligands, we noticed that all the QDs accorded a favorable effect on the transactivation activity and positively elevated PPAR-γ activation (Figure 2 (C-(a)). In comparison to the effects of RSG, QUENH showed the maximum levels of this protein at transcription level. This indicates it is acting as full agonist for the receptor. Also, QUEINH manifested almost similar level of expression to that of RSG. As expected, changes in receptor localization were prevented slightly by quercetin and translocated PPAR-γ lower than RSG. Although QUETSC accounted a good PPAR-γ expression at the protein level but relatively a slight attenuation of transcription factor binding was noted. Among the five QDs, only QUE2FH and QUESH behaved as partial agonists for PPAR-γ and significantly led to transactivation with a binding affinity lower than that of RSG and higher than quercetin.

A comparative assay was conducted with TGF-β1 to determine the basis for trans-location of PPAR-γ in its presence. Our analysis identified that the transactivation of PPAR-γ to nucleus was greatly reversed and reduced by TGF-β1 (Figure 2 (C-(b)). Whilst on administration of QDs to the cells, the transactivational binding was found to be in-creased and hence, elevated the translocation of PPAR-γ to nucleus to three-fold. Notably, with the treatment with TGF-β1, a maximum elevation for PPAR-γ protein translocation was sighted with quercetin. As a result, its derivatives exerted favorable effect on the transactivation activity. Among the QDs, QUETSC and QUEINH exhibited a slightly higher PPAR-γ binding activity than RSG at transcription level, while QUENH seems to have the lowest. Hence, QUENH yet again proved to be entirely a full agonist. As obvious, RSG showed feeble interaction which is in concordance with the above-mentioned protein assay results. Merely, even after the administration of TGF-β1, QUE2FH and QUESH enhanced the transactivation capability of PPAR-γ at moderate level and served as partial agonists. To this end, it is clear that the effect of QDs for PPAR-γ translocation was elevated even after the incorporation of TGF-β1, suggesting their role in the gene transcription by PPAR-γ.

### 5. TGF-β1-induced EMT markers was significantly affected by PPAR-γ partial activation

Previous reports suggest that PPAR-γ ligands may block TGF-β1-induced EMT [5, 19, 21]. Hence, we assessed the anti-metastatic effect of PPAR-γ partial agonists (QDs) on EMT process induced by TGF-β1. As expected, TGF-β1 treatment completely suppressed the E-cadherin expression. Comparatively, QDs significantly rescued TGF-β1–induced E-cadherin suppression at both the time intervals (Figure 3 (C)). As noted, changes in the EMT markers were greatly seen at 72h incubation. This is in accordance of the above results and indicates that the acquisition of EMT markers induced by TGF-β1 is time-dependent. According to the analysis, TGF-β1+QUENH, TGF-β1+QUE2FH and TGF-β1+QUESH manifested one-fold increase in E-cadherin expression compared to the control cells (only TGF-β1). While there was no significant difference on treating with other two QDs; TGF-β1+QUETSC and TGF-β1+QUEINH. This attributes that probably partial activation of PPAR-γ is able to prevent the complete loss of epithelial marker via these three QDs. Although quercetin elevated the protein expression, but RSG was unable to do so. In terms of mesenchymal markers, a complete reduction in the protein expression of Snail, Slug, Vimentin and Zeb-1 were perceived on TGF-β1 stimulation by all the five QDs at 72h compared to 24h. Conversely, no such effect was observed for N-cadherin (Figure 3(E)). These finding firmly proved that PPAR-γ may also play a critical role in modulating TGF-β1-induced EMT as well.

**Figure 3.**
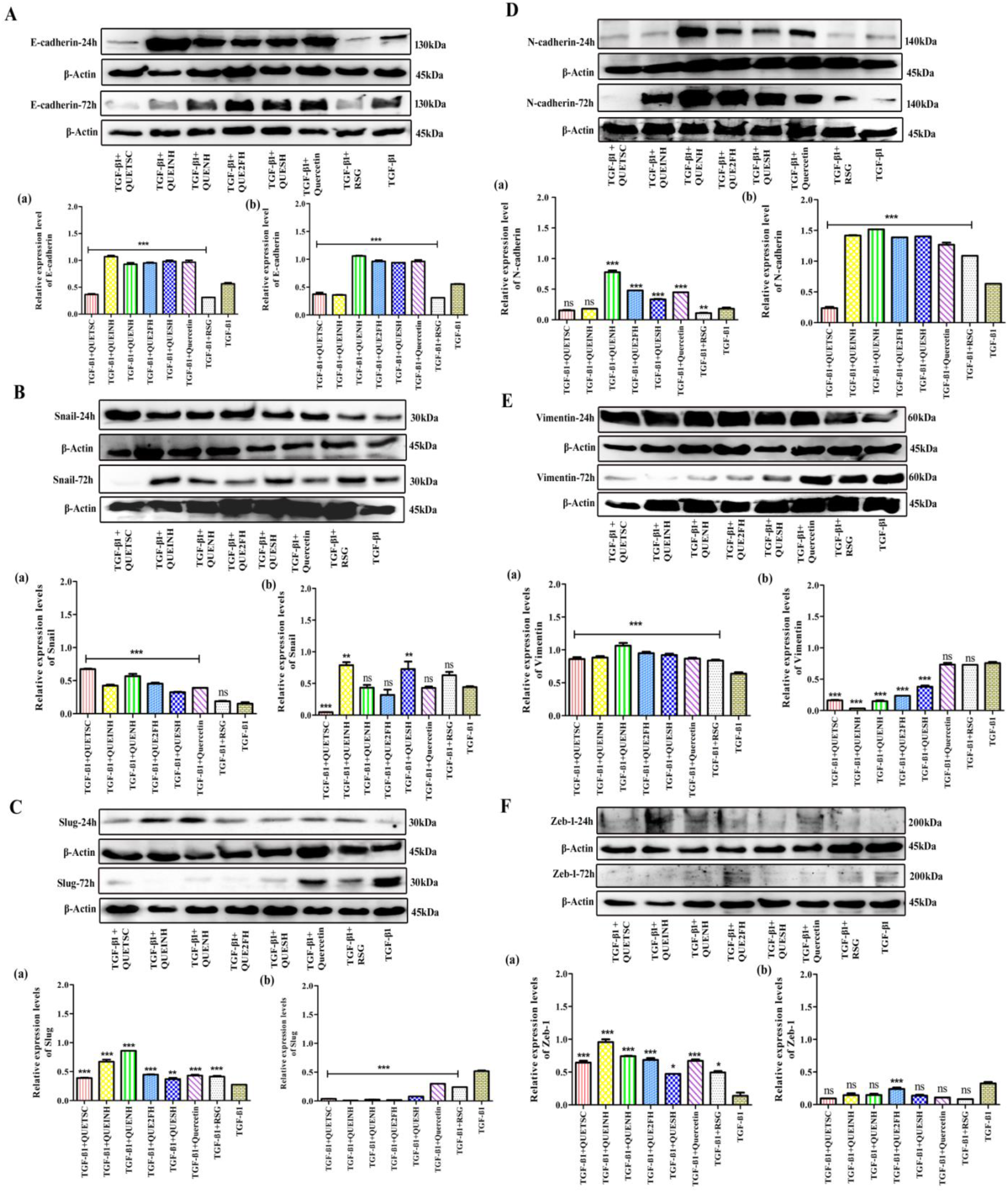
Effect of PPAR-γ ligands (synthesized QDs) on the acquisition of EMT markers on A549 cells induced with TGF-β1. Representative western blot images with their relative protein expression level of (A) E-cadherin (a)-24h and (b)-72h; (B) Snail (a)-24h and (b)-72h; (C) Slug (a)-24h and (b)-72h; (D) N-cadherin (a)-24h and (b)-72h; (E) Vimentin (a)-24h and (b)72h and (F) Zeb-1(a)-24h and (b)-72h with respect to β-actin. Each value presented as mean±standard deviation (SD) for three replicate determinations for each treatment (p<0.05, one-way ANOVA). The error bar represents SD; *p<0.05, **p<0.01 and ***p < 0.001 compared with control group-TGF-β1.

As QUETSC, QUE2FH, and QUESH served as partial agonists for PPAR-γ, thereafter we conducted a RT-PCR analysis to determine the EMT expression at mRNA level. The outcomes were observed to be in consistent with the RT-PCR analysis where the acquisition of mesenchymal markers; Slug, Snail and Vimentin were diminished with the exposure of three partial agonists (Figure S2). Although QUETSC downregulated the mesenchymal markers but was unable to upregulate the E-cadherin expression, therefore, it is certain that the effects were less pronounced. Thus, its partial activation process is questionable. In sum, the data implicated that amongst the three partial agonists, QUE2FH and QUESH exerted better effectiveness towards these markers and ought to be the most potential candidates to modulate the biochemical markers of EMT via PPAR-γ partial activation.

### 6. PPAR-γ partial agonists impeded cell migration in A549 cells upon TGF-β1 induction

TGF-β1 is highly known to contribute to the migration potential of cells to EMT process which is an indistinctive attribute. We investigated the functional consequence of partial activation of PPAR-γ on TGF-β1–induced A549 cell migration. It was observed that treatment of QDs remarkably reduced TGF-β1–induced migration (Figure 4 (B, C, D)) in A549 cells. The attenuation of tumor cell migration corresponds with the ability of partial agonist to inhibit mesenchymal markers. In contrast, RSG had no effect on the migration whilst quercetin was comparatively better than the full agonist. The result suggests that the effects of quercetin and RSG are specific to TGF-β1–induced responses. Comparatively, the migration activity of the cells was largely abolished by QUETSC. But due to its mere observation with EMT markers, the compound is not so efficient. We also assessed the morphological appearance of the treated cells with QDs, quercetin and RSG with or without TGF-β1 in order to evaluate the influence of these compounds on TGF-β1-induced EMT. Interestingly, cells induced with TGF-β1 in the presence of quercetin and RSG seem to undergo a slight change in morphology and tend to get scattered with the minute loss of cell-to-cell contacts (Figure 4 (A)). These findings proved the potency of QUE2FH and QUESH partial agonists in attenuating the cell migration which is well known for promoting tumorgenesis and metastasis.

**Figure 4:**
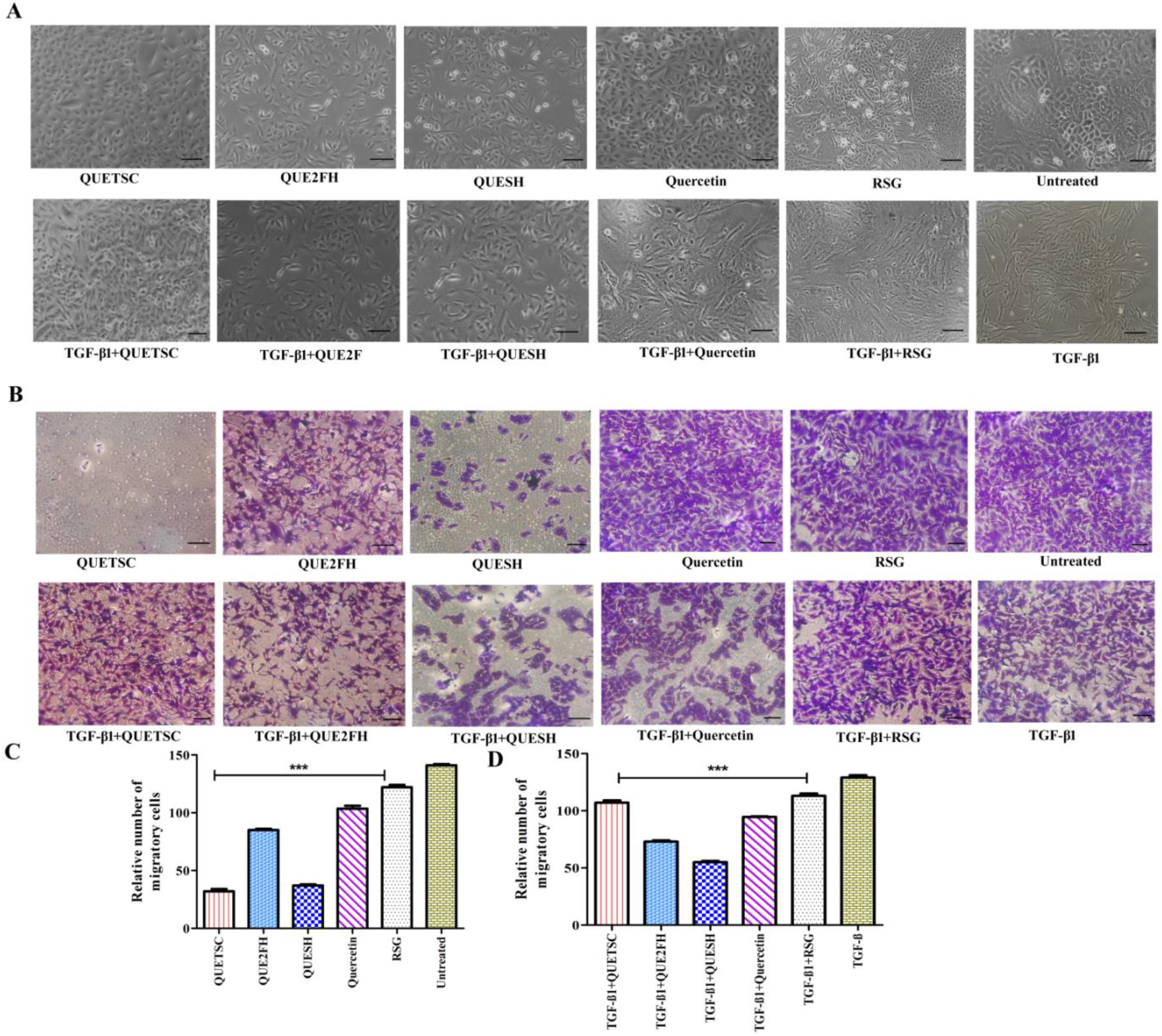
PPAR-γ partial agonists impeded cell migration in A549 cells upon TGF-β1 induction. (A) Representative Images of A549 cells at two timelines; 24h and 72h treated with IC_5_ doses of QUETSC, QUE2FH, QUESH, quercetin and RSG with or without TGF-β1 (5 ng/mL). (B) Representative Images of Transwell migration assay using Transwell insert. (A & B) The images were captured under Olympus microscope equipped with camera (Magnus Analytical MagVision, Version: x86). (C & D) Comparison of the number of migrating cells after treatment. Each value presented as mean±SD for three replicate determinations for each treatment (p<0.05, one-way ANOVA). The error bar represents SD; *p<0.05, **p<0.01 and ***p < 0.001 compared with control group-Untreated (C) &TGF-β1 (D). Scale bar: 100μm.

## Discussion

The present research extrapolates the ability of partial activation of PPAR-γ by novel synthesized derivatives of quercetin in modulating TGF-β1-induced EMT in lung cancer cells. Of the three PPAR-γ partial agonists tested, QUESH was the most potent modulator for EMT process. EMT is a well-studied mechanism that endows the cancer cells with increased resistance to apoptosis, elevated migratory and invasive capabilities, and thereafter increased aggressiveness. It is speculated that TGF-β1 influences these factors of EMT and serves as potential inducer for the progression of cancer cells. In fact, evidence shows that TGF-β1 is greatly expressed in lung cancer cells and promotes EMT [20-21]. During the process of EMT, TGF-β1 highly reprograms the epithelial cells to lose their properties and changes to mobile mesenchymal-like phenotype. This transition is well-reported by Reka et al wherein they observed A549 cells to undergo changes in their shape from cobble-shaped epithelial to elongated fibroblastoid form of mesenchymal characteristics in 72h induction of TGF-β1[3]. Consistent with most previous studies, our study showed that TGF-β1-induced (5 ng/ml) A549 cells underwent morphological difference clearly in 72h incubation.

As TGF-β1 is a crucial player to have a greater influence on EMT and having verified with PPAR-γ activation by QDs, here we extended our work to investigate the interplay between PPAR-γ, TGF-β1and synthesized QDs. Based on our protein analysis, the expression of PPAR-γ was seen to decelerate on TGF-β1 treatment. A report by Ramirez et al in the context of TGF-β1 influencing PPAR-γ showed an early and inductive effect of TGF-β1onPPAR-γ gene expression and hence led to an inverse in its action [14]. The authors reasoned out that TGF-β1can itself control PPAR-γ expression directly in lung cancer via mRNA expression in primary lung fibroblasts which tends to blunt its action. Moreover, the underlying mechanism responsible for such an effect is unknown, but the balance between TGF-β1 and PPAR-γ is likely to be determined by its gene expression. Further, we investigated how this effect is changed when PPAR-γ ligands binds to it. Our study showed that four out of five QDs, QUEINH, QUENH, QUE2FH and QUESH increased the PPAR-γ protein expression significantly after incorporation of TGF-β1. This suggested the QDs could inhibit the expression of TGF-β1and activated PPAR-γ in its presence. The reason can be ascribed to the phosphorylation of PPAR-γ which aids in attenuation of TGF-β1 on binding to PPAR-γ ligand [22]. PPAR-γ phosphorylation is said to be affected by various kinases such as Protein Kinase A and C (PKA, PKC), mitogen-activated protein kinases (ERK- and p38-MAPK), AMP Kinase (AMPK) and glycogen synthase kinase-3 (GSK3) [23]. Studies have demonstrated that these kinases tend to decrease the PPAR-γ phosphorylation which eventually leads to an increased PPAR-γ expression on binding of a ligand [24]. Activation of PPAR-γ is an important transcriptional event whereby a low transactivation potential leads to weak stimulation whilst high transactivation induces full activation [25].Talking of the partial activation, QUE2FH and QUESH behaved as partial agonists and reversed PPAR-γ protein expression in desired manner even after inducing TGF-β1 i.e., lower than quercetin and higher than RSG.

Since binding of ligand translocate PPAR-γ from cytoplasm to nucleus, we examined the effect of QDs on PPAR-γ transcriptional activity profile with or without TGF-β1. Moreover in the absence of TGF-β1, a similar trend was observed for PPAR-γ translocation to nucleus by QDs with that of our previous work wherein QUE2FH and QUESH behaved as partial agonists. While in the presence of TGF-β1, these two QDs enhanced the transactivation capability of PPAR-γ and adhered to partial activation phenomenon. In contrast, the synthetic PPAR-γ agonist, RSG triggered PPAR-γ to least extent and could not elevate its expression, thus indicating that RSG has negligible effect on translocation of PPAR-γ to nucleus. This which might be due to its strong ligand inducible properties and in the presence of TGF-β1 its effect got diminished [26]. These findings authenticate the behavioural pattern of QUE2FH and QUESH in accordance with partial activation. As we know the fact that TGF-β1 supresses PPAR-γ action at transcriptional level, a study conducted by Lakshmi et al proved that the recruitment Suppressor of Mothers against Decapentaplegic (SMAD) mediators are responsible for the effect [27]. The authors evidenced that the co-transfection of SMAD3/4 minimizes PPAR-γ promoter activity and thus induction of TGF-β1 downregulates PPAR-γ, however, the exact mechanism remain poorly understood [28-29].

Further to address the anti-metastatic effects of PPAR-γ ligands in TGF-β1-induced EMT, we explored the protein expression of various EMT markers. Interestingly, the QDs effectively reversed the acquisition of mesenchymal properties and suppressed their expression in TGF-β-stimulated lung cancer cells. These results were confirmed by Western blot, wherein Snail, Slug, Vimentin and Zeb-1 were observed to be downregulated and E-cadherin was upregulated by all the QDs. Most importantly, the full agonist, RSG had no effect on E-cadherin expression and was unable to upregulate. This is line with our transcriptional binding assay results in which the action of RSG was found to be reduced in the presence of TGF-β1. During the course of EMT in lung cancer, the cadherin switch is regarded as an important hallmark of metastasis and invasiveness [30]. The loss of E-cadherin in NSCLC is characterized by loss of cell-cell adhesionwhich eventually leads to cell mobility and invasion due to various transcriptional repressors such as Snail, Slug, Vimentin and Zeb-1. The zinc finger factors, Snail and Slug are known to be overexpressed in epithelial cell in lung cancer. The two factors are reported to enhance E-box interaction, ZEB1 and supresses E-cadherin [31]. A major observation was noted in the case of QUETSC as the compound did not fall into the hypothesis of the study. The compound failed to activate PPAR-γ in the presence of TGF-β1 at both transcriptional and protein level thereby could not elevate E-cadherin expression. However, our understanding of how TGF-β1 is affecting the action of QUETSC in lung cancer cell is still limited. Furthermore, the functional consequences of EMT such as cellular morphology and migration were observed to be blocked by PPAR-γ partial agonists; QUE2FH and QUESH. The crosstalk of PPAR-γ activation has been significantly shown to retard TGF-β1 signaling in other biological contexts as well with the participation of various ligands [24,32]. A quinolone alkaloid, Evodiamine was observed to counteract TGF-β1-induced EMT via PPAR-γ activation and Smad 2 signal pathway [32]. One possibility for the underlying molecular signaling mechanism for TGF-β1–induced EMT is the crucial role of SMAD mediators. SMAD phosphorylation is regulated by the stimulation of TGF-β1, wherein the Smad3 interacts with Smad4 by fundamental signaling kinases [33]. Numerous studies have indicated the suppression of EMT, migration and invasion induced by TGF-β1 by PPAR-γ activation with subsequent antagonizing the Smad3 signaling pathway [34]. However, the concept of partial activation have not been yet researched and elucidated for TGF-β1-induced EMT. Thusly, our study provides evidence to modulate EMT and migration in NSCLC via partial activation of PPAR-γ. The study is worthwhile to test and compare various other known PPAR-γ ligands and their effects on metastasis to make a significant impact on managing metastatic disease.

## Conclusion

In conclusion, for the first time, our study provided evidence for a protective role of novel synthesized derivatives of quercetin in counteracting TGF-β1-induced EMT in A549 cells via PPAR-γ dependent pathway. In light of evidences, the PPAR-γ stand out to be an attractive target as all the synthesized ligands induced anti-tumorigenic effects on A549 cells with its activation. Owing to their drug-like properties, the derivatives exhibited desired effects on inhibiting growth, EMT and colony formation of A549 cells at nanomolar concentrations. Of the five derivatives we screened, QUETSC, QUE2FH and QUESH was found to activate the protein cooperatively with a significant lower intrinsic activity than that of the full agonist RSG and higher activity over the weak agonist quercetin. Based on their putative binding modes, QUESH have been shown to impart to the high-affinity binding towards PPAR-γ receptor. Following PPAR-γ partial activation, QUESH ought to be the most potential to modulate the biochemical markers of EMT-induced by TGF-β1 with the inhibition of colony formation and migration invasiveness.These results suggest that QUESH appears as a promising PPAR-γ partial agonist which can be used as anti-metastatic agent.

## Acknowledgments

The authors are thankful to Dr. D. Y. Patil Biotechnology and Bioinformatics Institute, Dr. D. Y. Patil Vidyapeeth, Pune for the physical infrastructure. The authors also acknowledge the Department of Science and Technology Science and Engineering Research Board (DST-SERB), Govt. of India, New Delhi, (File Number: ECR/2016/000943) for financial support and utilizing an optimized supercomputer for dynamics calculations (File Number: YSS/2015/002035). S. Ballav is thankful to DST-SERB for Junior Research Fellowship.

## REFERENCES

1. Yang, J., Antin, P., Berx, G., Blanpain, C., Brabletz, T., Bronner, M., Campbell, K., Cano, A., Casanova, J., Christofori, G., Dedhar, S., Derynck, R., Ford, H. L., Fuxe, J., García de Herreros, A., Goodall, G. J., Hadjantonakis, A. K., Huang, R. Y. J., Kalcheim, C., Kalluri, R., … EMT International Association (TEMTIA) (2020). Guidelines and definitions for research on epithelial-mesenchymal transition. Nature reviews. Molecular cell biology, 21(6), 341–352. https://doi.org/10.1038/s41580-020-0237-9

2. Hao, Y., Baker, D., & Ten Dijke, P. (2019). TGF-β-Mediated Epithelial-Mesenchymal Transition and Cancer Metastasis. International journal of molecular sciences, 20(11), 2767. https://doi.org/10.3390/ijms20112767

3. Reka, A. K., Kurapati, H., Narala, V. R., Bommer, G., Chen, J., Standiford, T. J., & Keshamouni, V. G. Peroxisome proliferator-activated receptor-gamma activation inhibits tumor metastasis by antagonizing Smad3-mediated epithelial-mesenchymal transition. Molecular cancer therapeutics. 2010, 9(12), 3221–3232. DOI:10.1158/1535-7163.MCT-10-0570.

4. Burgess, H. A., Daugherty, L. E., Thatcher, T. H., Lakatos, H. F., Ray, D. M., Redonnet, M., Phipps, R. P., & Sime, P. J. (2005). PPARgamma agonists inhibit TGF-beta induced pulmonary myofibroblast differentiation and collagen production: implications for therapy of lung fibrosis. American journal of physiology.Lung cellular and molecular physiology, 288(6), L1146–L1153. https://doi.org/10.1152/ajplung.00383.2004.

5. Tan, X., Dagher, H., Hutton, C. A., & Bourke, J. E. (2010). Effects of PPAR gamma ligands on TGF-beta1-induced epithelial-mesenchymal transition in alveolar epithelial cells. Respiratory research, 11(1), 21. https://doi.org/10.1186/1465-9921-11-21.

6. Miyazono K. (2009). Transforming growth factor-beta signaling in epithelial-mesenchymal transition and progression of cancer. Proceedings of the Japan Academy. Series B, Physical and biological sciences, 85(8), 314–323. https://doi.org/10.2183/pjab.85.314

7. Choi S.S., Park J., Choi J.H. (2014) Revisiting PPAR-γ as a target for the treatment of metabolic disorders. BMB Rep.;47(11):599–608.

8. Taraghijou P.,Safaeiyan A.,Mobasseri M. andOstadrahimi A. (2012) The effect of n-3 long chain fatty acids supplementation on plasma peroxisome proliferator activated receptor gamma and thyroid hormones in obesity. J Res Med Sci.;17(10): 942–946.

9. Han, E., Jang, S.-Y., Kim, G., Lee, Y., Choe, E. Y., Nam, C. M., & Kang, E. S. (2016) Rosiglitazone Use and the Risk of Bladder Cancer in Patients With Type 2 Diabetes. Medicine;95(6): e2786.

10. Nesto, R. W., Bell, D., Bonow, R. O., Fonseca, V., Grundy, S. M., Horton, E. S., Le Winter, M., Porte, D., Semenkovich, C. F., Smith, S., Young, L. H., & Kahn, R. (2004). Thiazolidinedione use, fluid retention, and congestive heart failure: a consensus statement from the American Heart Association and American Diabetes Association. Diabetes care, 27(1), 256–263. https://doi.org/10.2337/diacare.27.1.256

11. Guan, Y., Hao, C., Cha, D. R., Rao, R., Lu, W., Kohan, D. E., Magnuson, M. A., Redha, R., Zhang, Y., & Breyer, M. D. (2005). Thiazolidinediones expand body fluid volume through PPARgamma stimulation of ENaC-mediated renal salt absorption. Nature medicine, 11(8), 861–866. https://doi.org/10.1038/nm1278

12. Da, C., Liu, Y., Zhan, Y., Liu, K., & Wang, R. (2016). Nobiletin inhibits epithelial-mesenchymal transition of human non-small cell lung cancer cells by antagonizing the TGF-β1/Smad3 signaling pathway. Oncology reports, 35(5), 2767–2774. https://doi.org/10.3892/or.2016.4661

13. Katsuno, Y., & Derynck, R. (2021). Epithelial plasticity, epithelial-mesenchymal transition, and the TGF-β family. Developmental cell, 56(6), 726–746. https://doi.org/10.1016/j.devcel.2021.02.028

14. Ramirez, A., Ballard, E. N., & Roman, J. (2012). TGFβ1 Controls PPARγ Expression, Transcriptional Potential, and Activity, in Part, through Smad3 Signaling in Murine Lung Fibroblasts. PPAR research, 2012, 375876. https://doi.org/10.1155/2012/375876

15. Camp, H. S., Li, O., Wise, S. C., Hong, Y. H., Frankowski, C. L., Shen, X., Vanbogelen, R., & Leff, T. (2000). Differential activation of peroxisome proliferator-activated receptor-gamma by troglitazone and rosiglitazone. Diabetes, 49(4), 539–547. https://doi.org/10.2337/diabetes.49.4.539

16. Gomez-Casal, R., Bhattacharya, C., Ganesh, N., Bailey, L., Basse, P., Gibson, M., Epperly, M., & Levina, V. (2013). Non-small cell lung cancer cells survived ionizing radiation treatment display cancer stem cell and epithelial-mesenchymal transition phenotypes. Molecular cancer, 12(1), 94. https://doi.org/10.1186/1476-4598-12-94

17. Gulbis, J. M., Kelman, Z., Hurwitz, J., O’Donnell, M., & Kuriyan, J. (1996). Structure of the C-terminal region of p21(WAF1/CIP1) complexed with human PCNA. Cell, 87(2), 297–306. https://doi.org/10.1016/s0092-8674(00)81347-1

18. Krishna, T. S., Kong, X. P., Gary, S., Burgers, P. M., & Kuriyan, J. (1994). Crystal structure of the eukaryotic DNA polymerase processivity factor PCNA. Cell, 79(7), 1233–1243. https://doi.org/10.1016/0092-8674(94)90014-0

19. Kao, H. F., Chang-Chien, P. W., Chang, W. T., Yeh, T. M., & Wang, J. Y. (2013). Propolis inhibits TGF-β1-induced epithelial-mesenchymal transition in human alveolar epithelial cells via PPARγ activation. International immunopharmacology, 15(3), 565–574. https://doi.org/10.1016/j.intimp.2012.12.018

20. Asselin-Paturel, C., Echchakir, H., Carayol, G., Gay, F., Opolon, P., Grunenwald, D., Chouaib, S., & Mami-Chouaib, F. (1998). Quantitative analysis of Th1, Th2 and TGF-beta1 cytokine expression in tumor, TIL and PBL of non-small cell lung cancer patients. International journal of cancer, 77(1), 7–12. https://doi.org/10.1002/(sici)1097-0215(19980703)77:1<7::aid-ijc2>3.0.co;2-y

21. Saji, H., Nakamura, H., Awut, I., Kawasaki, N., Hagiwara, M., Ogata, A., Hosaka, M., Saijo, T., Kato, Y., & Kato, H. (2003). Significance of expression of TGF-beta in pulmonary metastasis in non-small cell lung cancer tissues. Annals of thoracic and cardiovascular surgery : official journal of the Association of Thoracic and Cardiovascular Surgeons of Asia, 9(5), 295–300.

22. Li, R., Wang, Y., Liu, Y., Chen, Q., Fu, W., Wang, H., Cai, H., Peng, W., & Zhang, X. (2013). Curcumin inhibits transforming growth factor-β1-induced EMT via PPARγ pathway, not Smad pathway in renal tubular epithelial cells. PloS one, 8(3), e58848. https://doi.org/10.1371/journal.pone.0058848

23. Burns, K. A., & Vanden Heuvel, J. P. (2007). Modulation of PPAR activity via phosphorylation. Biochimica et biophysica acta, 1771(8), 952–960. https://doi.org/10.1016/j.bbalip.2007.04.018

24. Li, R., Wang, Y., Liu, Y., Chen, Q., Fu, W., Wang, H., Cai, H., Peng, W., & Zhang, X. (2013). Curcumin inhibits transforming growth factor-β1-induced EMT via PPARγ pathway, not Smad pathway in renal tubular epithelial cells. PloS one, 8(3), e58848. https://doi.org/10.1371/journal.pone.0058848

25. Capelli, D., Cerchia, C., Montanari, R., Loiodice, F., Tortorella, P., Laghezza, A., Cervoni, L., Pochetti, G., & Lavecchia, A. (2016). Structural basis for PPAR partial or full activation revealed by a novel ligand binding mode. Scientific reports, 6, 34792. https://doi.org/10.1038/srep34792

26. Fu, M., Zhang, J., Lin, Y., Zhu, X., Zhao, L., Ahmad, M., Ehrengruber, M. U., & Chen, Y. E. (2003). Early stimulation and late inhibition of peroxisome proliferator-activated receptor gamma (PPAR gamma) gene expression by transforming growth factor beta in human aortic smooth muscle cells: role of early growth-response factor-1 (Egr-1), activator protein 1 (AP1) and Smads. The Biochemical journal, 370(Pt 3), 1019–1025. https://doi.org/10.1042/BJ20021503

27. Lakshmi, S. P., Reddy, A. T., & Reddy, R. C. (2017). Transforming growth factor β suppresses peroxisome proliferator-activated receptor γ expression via both SMAD binding and novel TGF-β inhibitory elements. The Biochemical journal, 474(9), 1531–1546. https://doi.org/10.1042/BCJ20160943

28. Fu, M., Zhang, J., Lin, Y., Zhu, X., Zhao, L., Ahmad, M., Ehrengruber, M. U., & Chen, Y. E. (2003). Early stimulation and late inhibition of peroxisome proliferator-activated receptor gamma (PPAR gamma) gene expression by transforming growth factor beta in human aortic smooth muscle cells: role of early growth-response factor-1 (Egr-1), activator protein 1 (AP1) and Smads. The Biochemical journal, 370(Pt 3), 1019–1025. https://doi.org/10.1042/BJ20021503

29. Wei, J., Ghosh, A. K., Sargent, J. L., Komura, K., Wu, M., Huang, Q. Q., Jain, M., Whitfield, M. L., Feghali-Bostwick, C., & Varga, J. (2010). PPARγ downregulation by TGFß in fibroblast and impaired expression and function in systemic sclerosis: a novel mechanism for progressive fibrogenesis. PloS one, 5(11), e13778. https://doi.org/10.1371/journal.pone.0013778

30. Shi, Y., Wu, H., Zhang, M., Ding, L., Meng, F., & Fan, X. (2013). Expression of the epithelial-mesenchymal transition-related proteins and their clinical significance in lung adenocarcinoma. Diagnostic pathology, 8, 89. https://doi.org/10.1186/1746-1596-8-89

31. Bolós, V., Peinado, H., Pérez-Moreno, M. A., Fraga, M. F., Esteller, M., & Cano, A. (2003). The transcription factor Slug represses E-cadherin expression and induces epithelial to mesenchymal transitions: a comparison with Snail and E47 repressors. Journal of cell science, 116(Pt 3), 499–511. https://doi.org/10.1242/jcs.00224

32. Wei, J., Li, Z., & Yuan, F. (2014). Evodiamine might inhibit TGF-beta1-induced epithelial-mesenchymal transition in NRK52E cells via Smad and PPAR-gamma pathway. Cell biology international, 38(7), 875–880. https://doi.org/10.1002/cbin.10270

33. Jo, E., Park, S. J., Choi, Y. S., Jeon, W. K., & Kim, B. C. (2015). Kaempferol Suppresses Transforming Growth Factor-β1-Induced Epithelial-to-Mesenchymal Transition and Migration of A549 Lung Cancer Cells by Inhibiting Akt1-Mediated Phosphorylation of Smad3 at Threonine-179. Neoplasia (New York, N.Y.), 17(7), 525–537. https://doi.org/10.1016/j.neo.2015.06.004

34. Da, C., Liu, Y., Zhan, Y., Liu, K., & Wang, R. (2016). Nobiletin inhibits epithelial-mesenchymal transition of human non-small cell lung cancer cells by antagonizing the TGF-β1/Smad3 signaling pathway. Oncology reports, 35(5), 2767–2774. https://doi.org/10.3892/or.2016.4661

