## Supplementary files for "Partial Activation of PPAR-γ by Synthesized Quercetin Derivatives Modulates TGF-β1-Induced EMT in Lung Cancer Cells"

1. **Nuclear translocation**

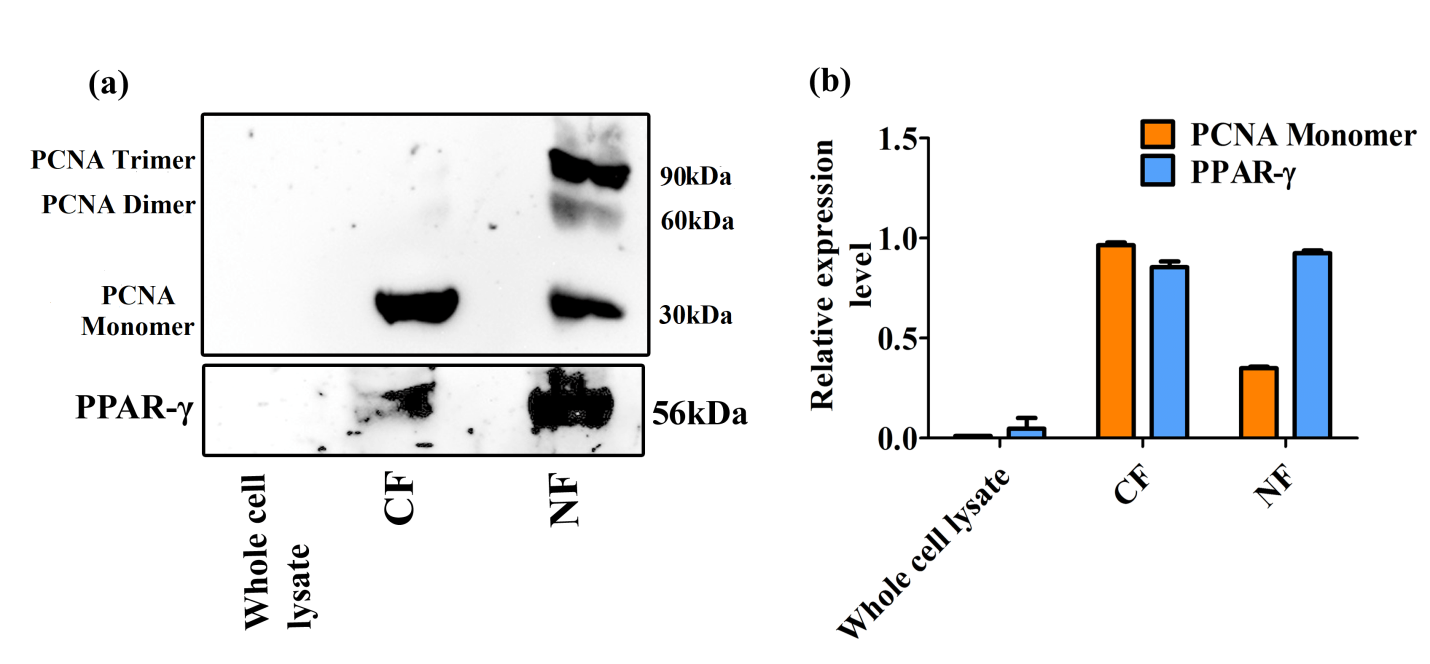

**Figure S1.** PPAR-γ was expressed in CF and NF followed by a graph representing the relative expression level for PCNA monomer and PPAR-γ in A549 cells.

2. **Quantitative Real-time (qRT-PCR) to determine the effect of PPAR-γ activation on EMT at mRNA level**

Total RNA was isolated from A549 cell lines treated with IC_5_ doses of QDs; QUETSC, QUE2FH and QUESH with quercetin as positive control with TGF-β1 (5ng/ml) for 24h by TRIzol solubilization and extraction method. Verso cDNA synthesis Kit (Thermo Scientific) provides robust transcription of RNA to create a complete cDNA pool. Real-time PCR was conducted using 2X Power SYBR™ Green PCR Master Mix (thermofischer, Cat. No.4367659). Each reaction was run in triplicate and contained 2µL of cDNA template along with 1.5µL of primers in a final reaction volume of 20µL. Cycling parameters were 95˚C for 2 min to activate DNA polymerase for 1 cycle, then 40 cycles of 95˚C for 2min, 58˚C for 30sec and 72˚C for 1 min. The following primer sequences were used for the reactions:

**GAPDH RT-PCR sequence: Forward Primer:**  AGACAGCCGCATCTTCTTGT, **Reverse Primer:** CTTGCCGTGGGTAGAGTCAT

**Human Vimentin RT-PCR sequence: Forward Primer:** TACAGGAAGCTGCTGGAAGG **Reverse Primer:** ACCAGAGGGAGTGAATCCAG

**Human Human Snail RT-PCR sequence:Forward Primer:** GAGGCGGTGGCAGACTAG **Reverse Primer:** GACACATCGGTCAGACCAG

**Human Human Slug RT-PCR sequence:Forward Primer:** CATGCCTGTCATACCACAAC **Reverse Primer:** GGTGTCAGATGGAGGAGGG

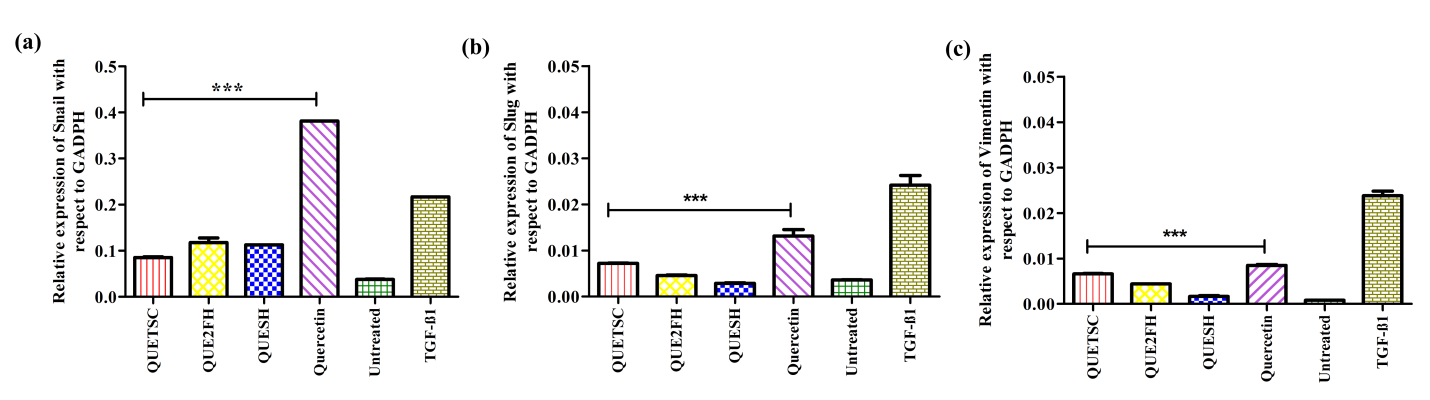

**Figure S2. PPAR-γ partial agonists (QUETSC, QUE2FH, QUESH) inhibits EMT in A549 cells induced by** **TGF-β1.** A549 cells were stimulated with TGF-β1 (5ng/ml) in the presence of treatment with IC_5_ doses were treated with QUETSC, QUEINH, QUESH and quercetin for 24h. Mesenchymal markers (a) Snail (b) Slug (c) Vimentin were monitored by RT-PCR. GADPH levels were assesed as loading control for whole cell-extracts. Each value presented as mean±SD for three replicate determinations for each treatment (p<0.05, one-way ANOVA).The error bar represents SD; *p<0.05, **p<0.01 and ***p < 0.001 compared with control group-TGF-β1.

3. **Western Blotting**

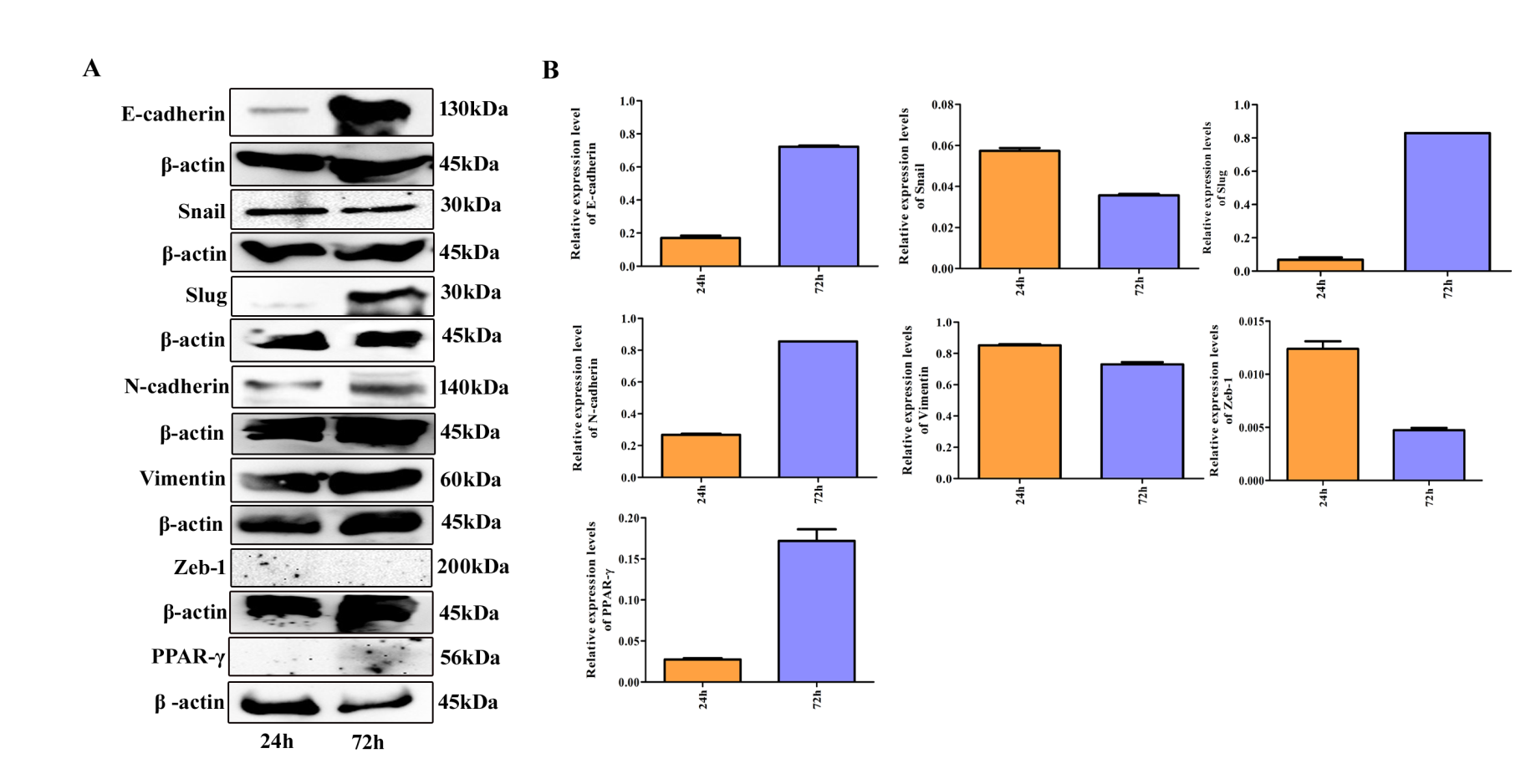

**Figure S3. Western blots for respective protein markers of A549 cells induced with TGF-β1 without treatment.** (A) A549 cells were treated with TGF-β1 (5 ng/mL) for 24h and 72h and proteins were obtained, separated on SDS–PAGE, probed with respective protein antibodies. (B) Relative protein expression level with respect to β-actin.
