## Supplementary figures and images for "Partial Activation of PPAR-γ by Synthesized Quercetin Derivatives Modulates TGF-β1-Induced EMT in Lung Cancer Cells"

### Supplementary Figure 1

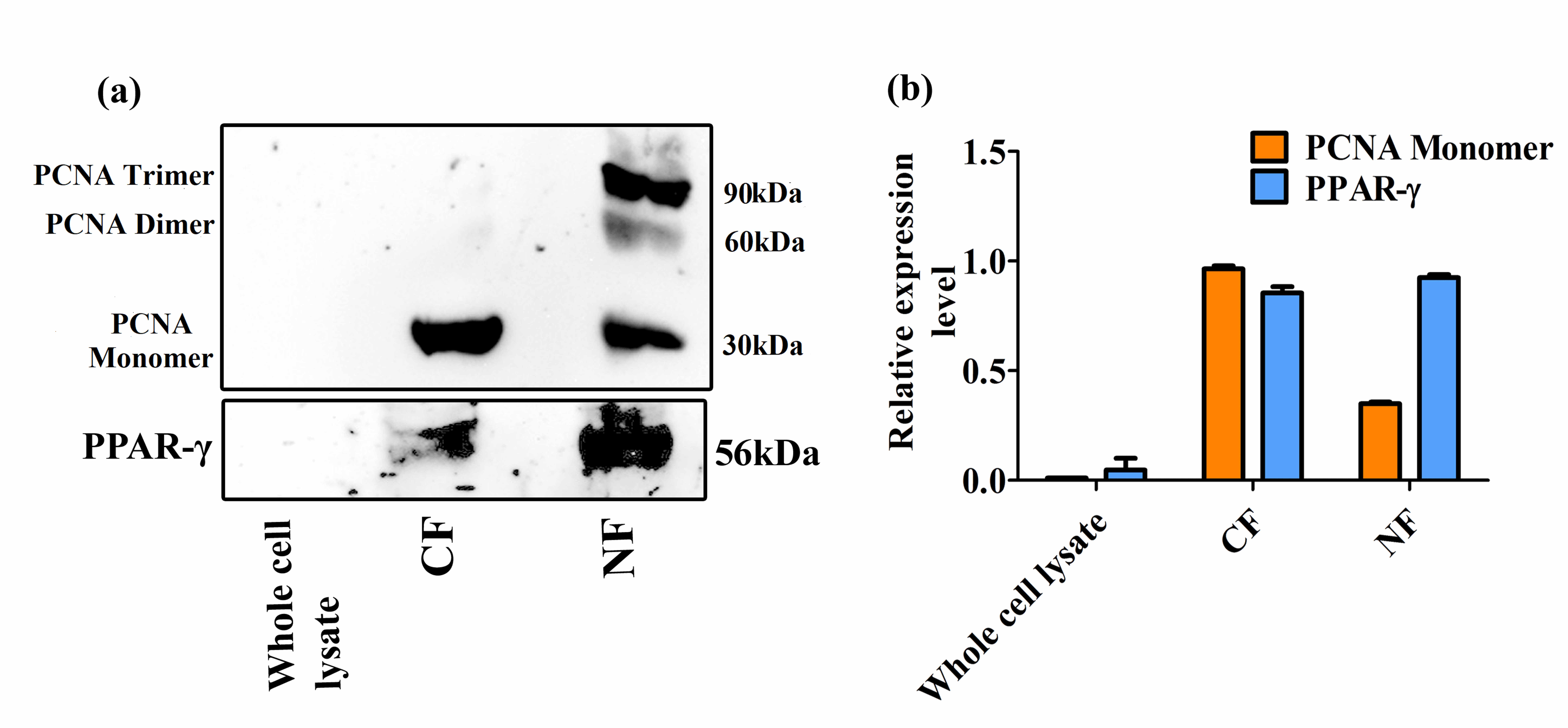

### Supplementary Figure 2

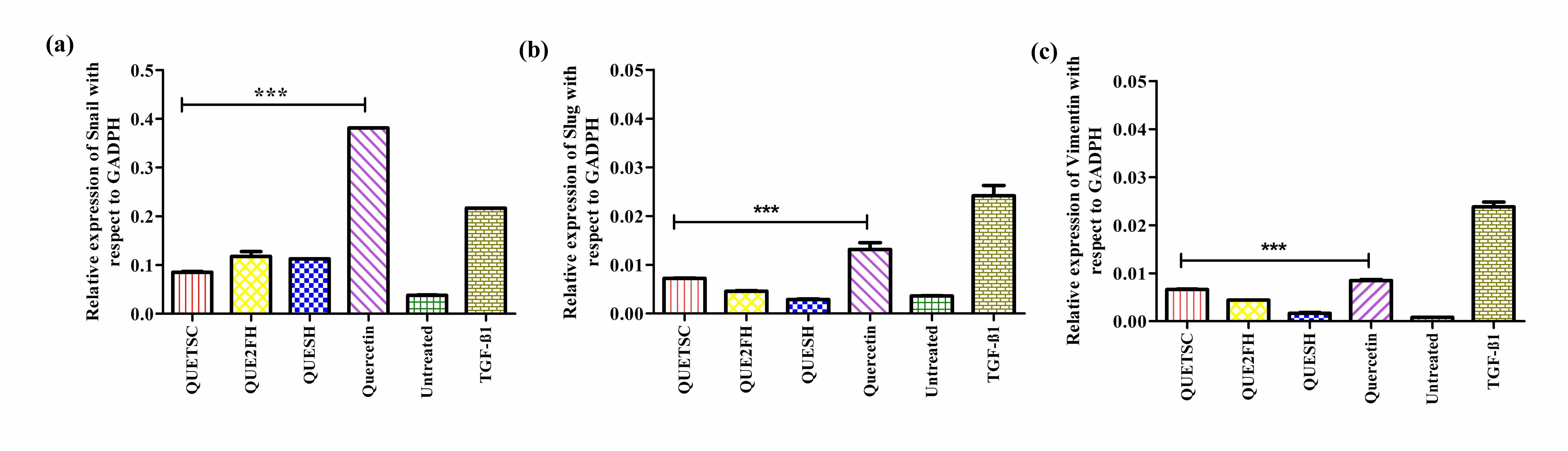

### Supplementary Figure 3

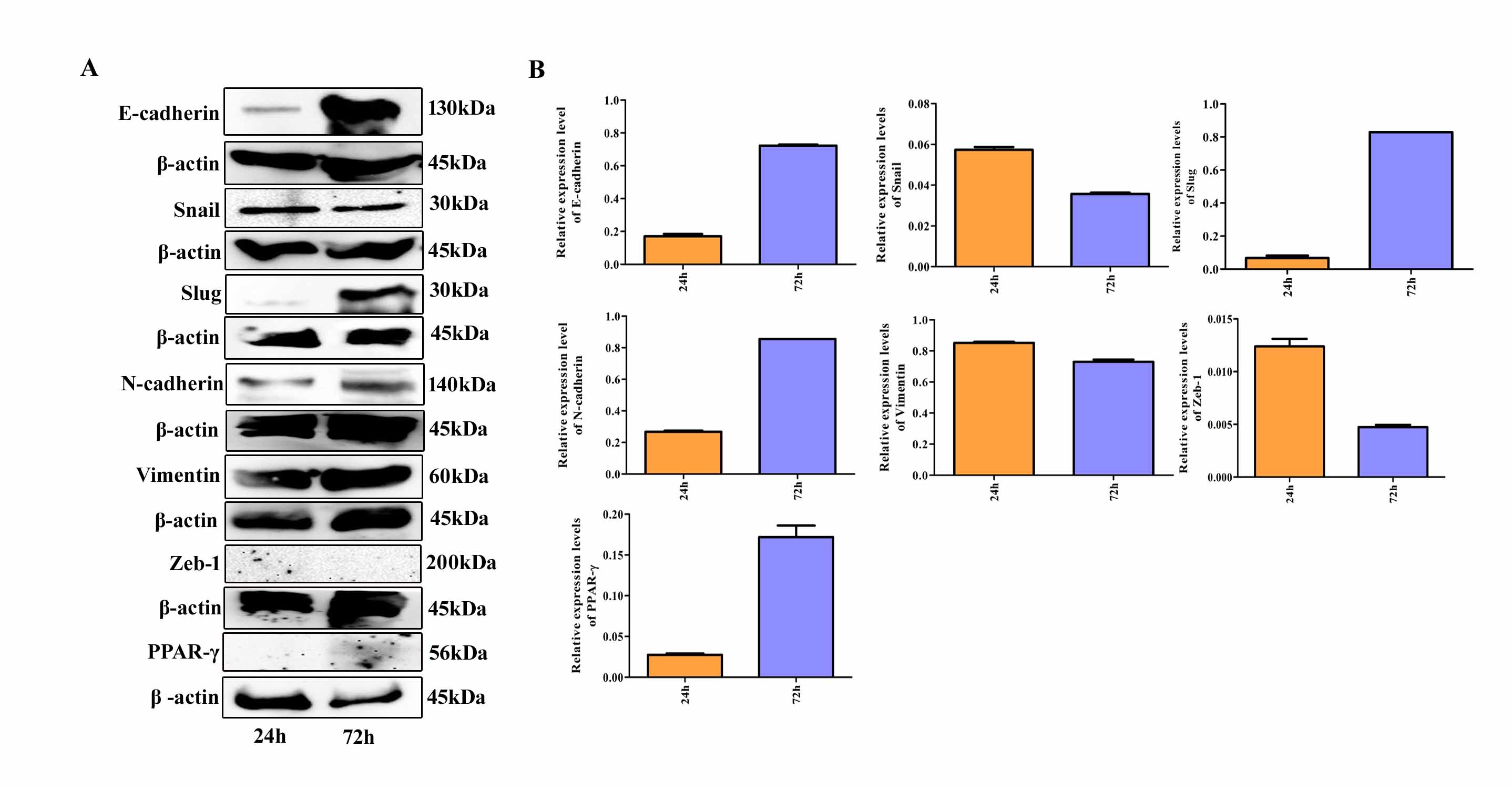
